# Anti-tumor activity of camptothecin analog conjugate of a RSPO4-based peptibody targeting LGR4/5/6 in preclinical models of colorectal cancer

**DOI:** 10.1101/2024.10.08.616548

**Authors:** Yukimatsu Toh, Ling Wu, Jianghua Tu, Zhengdong Liang, Adela M. Aldana, Jake J. Wen, Li Li, Sheng Pan, Rowe H. Julie, Martha E. Hensel, Carolyn L. Hodo, Rick A. Finch, Kendra S. Carmon, Qingyun J. Liu

## Abstract

Antibody-drug conjugates (ADCs) have emerged as a major modality of targeted cancer therapy, yet no ADC has been approved specifically for colorectal cancer (CRC). LGR4/5/6 (leucine-rich repeat containing, G protein-coupled receptor 4, 5, 6) are three related receptors that are expressed at high levels together or alternately in nearly all cases of CRC. ADCs targeting LGR5 have been shown to have robust anti-tumor potency, but not all CRC cells express LGR5 and LGR5-positive tumor cells may lose LGR5 expression due to cancer cell plasticity. R-spondin 4 (RSPO4) is a natural protein ligand of LGR4/5/6 with high affinity for all three receptors. We fused a mutant form of RSPO4 furin domain that retains high affinity binding to LGR4/5/6 to human IgG1 Fc to create a peptibody designated R462. Conjugation of R462 with a camptothecin analog designated CPT2 at eight drugs per peptibody led to the generation of R462-CPT2 that showed highly potent cytotoxic activity in vitro in CRC cell lines expressing any of LG4/5/6. In cell line xenograft and PDX models of CRC, R462-CPT2 demonstrated robust anti-tumor effect. Importantly, R462-CPT2 showed no major adverse effect at therapeutically effective dose levels. These results strongly support the use of RSPO ligand drug-conjugates that target LGR4/5/6 simultaneously for the treatment of CRC.

## Introduction

Colorectal cancer (CRC) remains a major cause of cancer-related deaths in the world, making it a critical area of research and development in oncology (1). Traditional chemotherapies are often accompanied by significant side effects and can be ineffective against advanced or metastatic disease (1,2). Immune checkpoint inhibitors are only effective in ∼6% of CRC patients, largely restricted to those with highly mutated tumors, such as those of microsatellite instable (MSI) nature(2). The Wnt/β-catenin signaling pathway plays a critical role in CRC tumorigenesis and progression with mutations in this pathway being found in the majority of CRC cases (3,4). Leucine-rich repeat containing, G protein-coupled receptors 4, 5, and 6 (LGR4/5/6) are three related membrane receptors that bind to R-spondins (RSPOs) with high affinity and potentiate Wnt signaling in response (5,6). In fact, gain-of-expression gene fusions of RSPO2 and RSPO3 were identified a small subset of CRC without mutations in APC or β-catenin genes (7). RSPOs comprise an N-terminal furin-like domain with two repeats (Fu1 and Fu2) and thrombospondin-like domain (TSP) at the C-terminus(8). The furin-like domain is both necessary and sufficient to bind LGRs to potentiate Wnt signaling whereas the TSP domain enhances the potency of furin-like domain (9–12). Crystal structure analysis revealed that the Fu1 domain binds to the E3 ubiquitin ligases (RNF43 and ZNRF3) while the Fu2 domain binds to the extracellular domain of LGR4/5 (13–16). Mechanistically, RSPO and LGR4 form a complex to inhibit the function of two E3 ligases (RNF43 and ZNRF3) that would otherwise ubiquitinate Wnt receptors for degradation, leading to higher Wnt receptor levels and stronger signaling (17–19). The RSPO-LGR5 complex, on the other hand, adopts a configuration that is unable to interact with the E3 ligases yet potentiates Wnt/β-catenin signaling through a different mechanism (20,21).

LGR4/5/6 are often co-upregulated in various cancer types, particularly in cancers of the gastrointestinal system, with LGR4 co-expressed with LGR5 or LGR6.(22–27) Furthermore, LGR5 has been shown to be enriched in CRC stem cells (28–31) and LGR5-positive cancer cells were shown to fuel the growth of primary tumors and metastasis (32,33). LGR6 is a marker of cancer stem cells of squamous cell carcinoma (31). The Cancer Genome Atlas’s (TCGA’s) RNA-Seq data of colorectum, liver, and stomach cancers confirmed that LGR4 expression at high levels in nearly all cases while LGR5 and LGR6 were co-expressed with LGR4 in the majority of CRC and substantial fractions of liver and stomach cancer. LGR4/5/6 were also found to be highly expressed in liver metastasis of CRC(34,35). These evidences support that targeting LGR4/5/6 together may provide an effective approach for the treatment of metastatic CRC.

Antibody-drug conjugates (ADCs) have become a major modality of cancer therapy (36,37). ADCs consist of a monoclonal antibody linked to a cytotoxic payload via a chemical linker (36,37). The antibody component specifically targets antigens overexpressed on cancer cells, delivering potent cytotoxic agent directly to the tumor, thereby reducing systemic toxicity.

Previously, we reported the use of a modified, antagonistic RSPO4 peptibody that was based on the fusion of a modified RSPO4 furin domain to human IgG1-Fc domain to deliver the cytotoxin monomethyl auristatin E (MMAE) into cancer cells expressing any of LGR4/5/6 (38), which is similar to the ADC approach except that the ligand-drug conjugate enables the targeting of LGR4/5/6 simultaneously. As microtubule inhibitors such as MMAE are not particularly effective against CRC (39), we generated RSPO4-furin peptibody conjugated with a derivative of camptothecin which is known to be particularly effective against CRC (2,40). Here we report the generation and characterization of R462-CPT2, an antagonistic RSPO4-furin-based peptibody-drug conjugate (PDC) that contained eight molecules of a camptothecin analog called CPT2 with improved potency and solubility using a chemoenzymatic conjugation method (41–43). R462-CPT2 showed highly potent cytotoxic activity across CRC cell lines expressing LGR4/5/6 at various levels in vitro and robust anti-tumor effect in cell line xenograft and PDX tumors of CRC in vivo without major adverse effect.

## Materials and Methods

### Peptibody and PDC preparation

The RSPO4-Fc fusion protein (R462) was designed based on the RSPO4 furin domain (aa 32-137), utilizing the following polypeptide chain: RSPO4 furin domain with Q65R fused to human IgG1-Fc, incorporating the N297Q mutation and connected by a G4S x3 linkers. Vectors for R462 were constructed using PCR-based In-Fusion cloning (Takara Bio). R462 was produced using the Expi293™ Expression System (Thermo Fisher Scientific) and purified using CaptivA® HF Protein A Resin (Repligen) in a gravity column, followed by a size-exclusion column of HiLoad 16/600 Superdex 200 pg (Cytiva). A solution of R462 (5 mg/mL in PBS (pH 7.2)) underwent treatment with 8% Activa® TI Transglutaminase (MTG, Ajinomoto) and 40 molar equivalents of N-(Amino-PEG2)-N-bis (PEG3-Azide) (BrodPharm) linker overnight at room temperature as part of the conjugation process. The excess linker and MTG were thoroughly removed by treating the product with protein A Resin, followed by 2 hours of vortexing at room temperature. Next, 1.5 molar equivalents of linker-drug (from a 20 mg/mL DBCO-PEG8-Val-Lys-Gly-14-aminomethy (CPT2) (Levena) in dimethyl sulfoxide (DMSO) stock solution) were added to the previous step production, while keeping the residual DMSO concentration below 10% (v/v). The mixture was incubated at room temperature for four hours, followed by purification over a size-exclusion column to remove excess reagents. In this step, the resulting PDC was also buffer exchanged into a formulation buffer (20 mM sodium succinate, 6% Trehalose, pH 5.0), and the final PDC was stored at -80 °C. R465-CPT2 was generated in the same scheme.

The proteins and the PDC samples were analyzed on an Agilent 6538 UHD Accurate-Mass Quadrupole Time-of -Flight (Q-TOF) LC/MS system coupled with an Agilent 1200 series HPLC. The LC separation was carried out on an Agilent PLRP-S reversed phase column (50 x 2.1 mm, 5µm, 1000Å). The solvent was 2 % acetonitrile, 97.9% water, and 0.1% formic acid (A); 80% acetonitrile, 19.9% water, and 0.1% formic acid (B). A gradient of 25% -90% B was applied over 20 min at a flow rate of 0.2 mL/min. The samples were injected after being diluted with 25% acetonitrile, 75% water, and 0.1% formic acid. The Q-TOF was operated in ESI positive mode with capillary voltage 3500 V, drying gas flow rate of 7L/min and the source temperature of 325 °C. Spectra were acquired in MS1 scan from 4.5 -20 min each run over the mass range of 650-2800 m/z. The raw data were processed using Agilent MassHunter BioConfirm software (Version B.04.00) which uses the Maximum Entropy deconvolution algorithm for the accurate molecular mass calculation, the deconvoluted mass range was set at 20,000 to 100,000 Daltons.

### Cell Lines and cell-based binding assays

All cancer cell lines were purchased from ATCC. HEK293 cells expressing LGR4, LGR5, or LGR6 were described previously reported (5,44). For whole-cell-based binding assays, stable HEK293T cells expressing LGR4 or LGR5 were seeded onto poly-D-lysine-coated 96-well plates at 50,000 cells/well and cultured overnight. Serial dilutions of protein samples and PDCs were added for 2 hours at 4 °C to prechilled cells. Plates were washed in PBS three times and fixed in 4% formalin for 30 min, and incubated with goat anti-human-Alexa-555 (Invitrogen) for 1 h at room temperature. Plates were washed with PBS and fluorescence intensity was quantified using a Tecan Infinite M1000 plate reader (excitation nm, emission nm). The data were analyzed using GraphPad Prism software, and the dissociation constant (Kd) values were determined.

### In vitro cytotoxicity assays

Cells were seeded to a 96-well plate at an appropriate density. Serial dilutions of PDCs were added to the cells, followed by incubation at 37 °C for 5 days to evaluate cell viability. Cell viability was assessed using the CellTiter-Glo® assay (Promega) for cell lines, following the manufacturer’s protocol. The luminescence was then measured using a Tecan Infinite M1000 plate reader. Data presented are representative results from a minimum of three independent experiments. IC50 values, compared with other controls, were determined using inhibition dose-response curve fitting in GraphPad Prism software.

### Pharmacokinetics of peptibody and PDC in mice

Animal studies were carried out in strict accordance with the recommendations of the Institutional Animal Care and Use Committee of the University of Texas at Houston (Protocol number AWC-22-0080,AWC-20-0144, and AWC-23-0106). R462 and R462-CPT2 were intraperitoneally injected at a dose of 5 mg/kg into female C57BL/6J mice aged 11 weeks, followed by blood sampling at intervals of 1, 24, 48, and 72 hours after the injection. The levels of PDC and the peptibody in the plasma were assessed through ligand-binding assays.

Specifically, the plasma samples were diluted 10-fold or 20-fold and then exposed to HEK293T-LGR5 cells. The concentrations of the ligands were deduced from standard curves generated using R462 with the identical cell type.

### Cell line and patient-derived xenograft studies

Cell line-derived xenograft tumors were established by injecting 5 million LoVo cells, mixed with Matrigel, subcutaneously into 9-week-old female nu/nu nude mice from Charles River Laboratories. Once the tumors reached an appropriate size, the tumor-bearing mice were randomized into 3 treatment groups of 6-7 mice each: vehicle (PBS), R462-CPT2, or R465-CPT2. Patient-derived xenograft models were established in NOD scid gamma (NSG) mice from Charles River. The CRC-001 (#TM00849) model was purchased from Jackson Laboratory as previously published (45), while XST-GI-007 and XST-GI-010 models were established at UTHealth with patient consent and approvals (HSC-MS-20-0327, HSC-MS-21-0074). Tumor fragments 2-3 mm in size were implanted subcutaneously into 6-8-week-old female mice. When tumors reached sufficient size, mice were randomized into treatment groups (N=4-6 per group) with equivalent average starting tumor volumes. Mice received intravenous injections of vehicle, R462-CPT2, or R465-CPT2 as indicated. PDCs were dosed intraperitoneally at 5 mg/kg every other day for 7 total doses. Both CDX and PDX model of tumor volumes were measured bi-weekly using: volume = (length x width^2)/2. Mice were euthanized when tumors reached 15 mm diameter. Tumor growth inhibition percentage was calculated as [1 - (tumor volume change in treatment / tumor volume change in vehicle)] x 100%.

### Toxicity study in mice

Toxicity study in mice (C57/Bl mice) were performed and R462-CPT2 was administered IP at doses of 0 (vehicle), 5, 10, and 15 mg/kg every other day for three doses total (3M/3F per group). The animals were monitored for general well-being and body weight. At terminated on Day 8, complete blood account and serum analysis were carried out and a total of 25 organs were collected for weights and histopathology analysis.

### Statistical analysis

Data analysis was performed using GraphPad Prism software. The data are presented as mean ± SEM or SD, as specified in the Results segment. For statistical testing of in vivo experiments with more than two groups, one-way ANOVA was applied along with Tukey’s multiple comparisons test. When comparing just two groups from these datasets or experiments, an unpaired, two-tailed Student’s t-test was utilized instead. A p-value less than 0.05 was considered statistically significant for all analyses.

## Results

### An RSPO4-furin peptibody was conjugated with eight molecules of CPT2 per peptibody

The Fu1 and Fu2 domain of RSPOs are structurally relatively independent and bind to the extracellular domains of the E3 ligases (RNF43/ZNRF3) and LGR4/5/6, respectively, as illustrated by a modeled LGR4-RSPO-ZNRF3 heterotrimer (Fig. 1a-b). Previously, we reported the generation and characterization of the peptibody R427 that comprised RSPO4 furin domain with a Q65R mutation in the Fu1 domain fused to the N-terminus of human IgG1-Fc that was tagged with LLQGA at the C-terminus of Fc. The LLQGA tag was used for direct conjugation of linker-payload by the microbial transglutaminase (46). R427 was able to bind LGR4/5/6 with high affinity without potentiating Wnt/β-catenin signaling, and even antagonized activity of wildtype RSPO1 and RSPO2 (38). However, R427 only contained two Gln (Q) residues in the LLQGA tag for conjugation, one on each Fc chain, and only four drugs could be conjugated even with a bis linker. Since camptothecin(CPT)-based payloads generally require eight molecules per antibody/peptibody for maximum efficacy (47), we created a peptibody called R462 that is similar to R427 except that the LLQGA tag was removed and N297 in the Fc domain was mutated to Q, creating two Gln residues (Q295 and Q297) that could be used by the microbial transglutaminase for conjugation (Fig. 1b) (43). We also created a control peptibody designated R465 that is identical to R462 except that it had two additional mutations (F99A and F103A) in the RSPO4-Fu2 domain to abolish its binding to LGR4/5/6 (Fig. 1c) (48).

**Figure 1.**
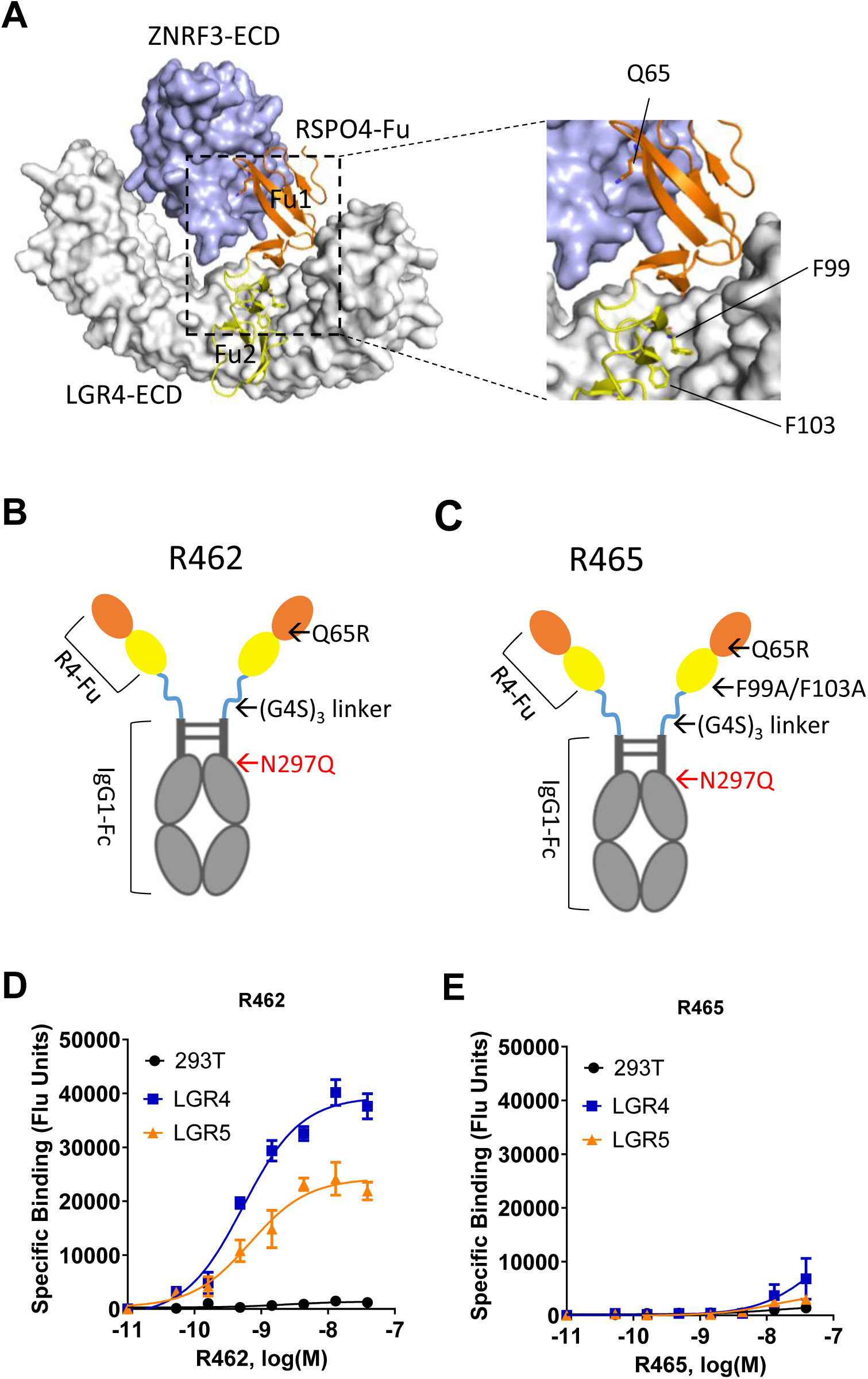
Design and characterization of peptibodies R462 and R465. **A**, A trimeric structure of RSPO4-Fu, ZNRF3-ECD, and LGR4-ECD modeled using the AlphaFold server. The left panel shows the overall structure. The right panel shows a close-up view of three key residues that are mutated to disrupt trimer formation. ZNRF3-ECD (light blue) and LGR4-ECD (white) are depicted as surfaces, while RSPO4-Fu (Fu1: orange; Fu2: yellow) is shown as a ribbon. The key residues are displayed as sticks. **B-C**, Schematic diagram of peptibody R462 (**B**) and R465 (**C**) showing the locations of amino acid residues that were changed for activity and conjugation. **D**, Saturation binding of R462 to HEK293T cell and HEK293T cells stably expressing LGR4 or LGR5. **E**, Saturation binding of R465 to HEK293T cells and HEK293T stably expressing LGR4 or LGR5. Error bars are S.E.M (n = 3).

R462 and R465 were expressed in Expi293™ cells and purified to apparent homogeneity (Supplementary Fig. S1). To assess the specificity and binding affinity of R462 and R465, saturation binding analysis were performed on HEK293T expressing LGR4 or LGR5 as previously described (38). R462 showed high affinity binding to both LGR4 and LGR5 with Kd of 0.5 nM and 0.7 nM respectively, while little binding to parental HEK293T cells was observed (Fig. 1d). In contrast, the control peptibody R465 demonstrated a near complete loss of binding to LGR4 and LGR5 as expected (Fig. 1e). These results demonstrated that R462 was able to bind to both LGR4 and LGR5 with similar affinity, and is expected to bind to LGR6 in a similar fashion as demonstrated with R427 (38).

Next, we conjugated a CPT derivative designated CPT2 to R462 and R465 using an chemoenzymatic conjugation method (41,42). CPT2 was shown to have higher potency in inhibiting Type I DNA topoisomerase with better solubility when compared to the two camptothecin derivates SN38 and deruxtecan that were approved in the FDA-approved ADCs (41). We had CPT2 with a PEG8-VKG linker capped with DBCO custom-synthesized (Levena BioPharma) to generate DBCO-PEG8-VKG-CPT2 (Fig. 2a). To reach a DAR (drug-to-antibody ratio) of 8, a bis linker (N-(amino-PEG2)-N-bis (PEG3-azide)) was first attached to R462 and R465 using microbial transglutaminase. Mass spectrometry analysis confirmed that all four Gln residues were conjugated (Fig. 2c). Next, DBCO-PEG8-VKG-CPT2 was attached to the azido groups using click chemistry (Fig. 2A) (42,49). The resulting peptibody drug conjugates (PDCs) R462-CPT2 and R465-CPT2 were purified by affinity chromatography followed by gel filtration to remove any aggregates or oligomers (Fig. 2b). Mass spectrometry analysis was conducted to evaluate the DAR of the conjugates. The results confirmed the successful attachment of eight CPT2 molecules to the Fc region of the PDCs (Fig. 2c). In the receptor binding assays, R462-CPT2 showed Kd of 2.4 and 1.3 nM on cells expressing LGR4 or LGR5, respectively (Fig. 2d). Interestingly, compared to unconjugated R462, R462-CPT2 showed low affinity but significant binding to vector control cells, suggesting that conjugation led to non-LGR4/5-dependend binding. Indeed, R465-CPT2 also showed a similar level of binding on vector control cells and LGR4/5 cells (Fig. 2e), suggesting that the conjugation led to a similar level of LGR4/5-independent binding. These findings suggest that the R462-CPT2 retained high affinity, specific binding to LGR4 and LGR5. Both R462-CPT2 and R465-CPT2 displayed a similar level of LGR4/5-independent low affinity binding.

**Figure 2.**
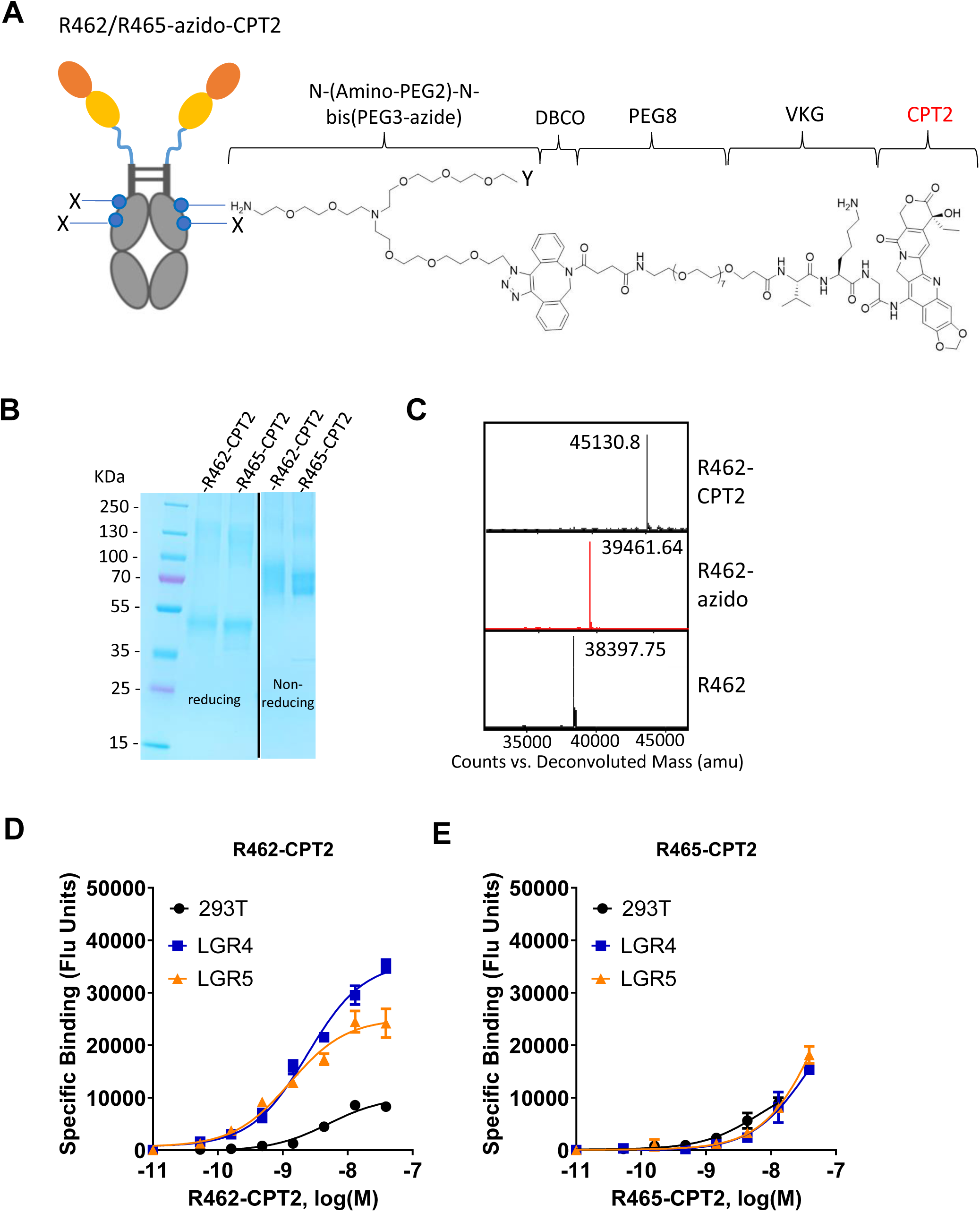
Design, conjugation and characterization of PDCs R462-CPT2 and R465-CPT2. **A**, Schematic diagram showing the structure of linker-payload conjugated to the peptibodies with click chemistry. Each peptibody has 8 molecules of linker-payloads conjugated. **B**, SDS-PAGE analysis of R462-CPT2 and R465-CPT2 under reducing (left side) and non-reducing condition. **C**, Mass spectra of R462, R462-azido, and R462-CPT2. **D**, Saturation binding of R462-CPT2 to HEK293T cells and HEK293T cells expressing LGR4 or LGR5. **E**, Saturation binding of R462-CPT2 to HEK293T cells and HEK293T cells expressing LGR4 or LGR5. Error bars are S.E.M (N = 3).

### R462-CPT2 showed highly potent activity in inhibiting the growth of CRC cell lines in vitro

To evaluate and compare in vitro cytotoxic activities of the PDCs (R462-CPT2 and R465-CPT2), we selected a panel of CRC cell lines expressing LGR4/5/6 receptors endogenously at various levels based on the Cancer Cell Line Encyclopedia’s (CCLE’s) RNA-seq data (Table 1).

**Table 1.**
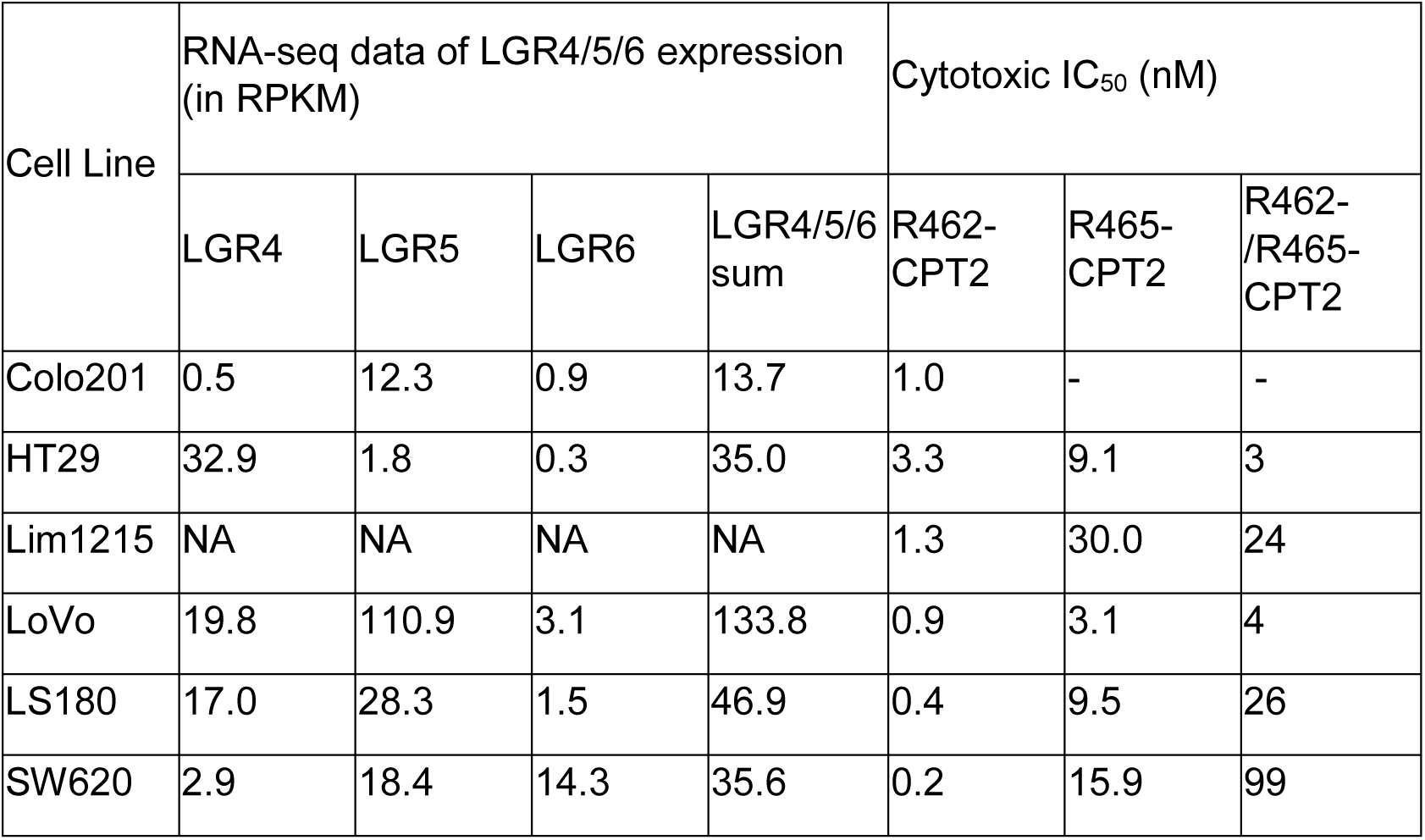
RNA-seq value (RPKM) of LGR4/5/6 and In vitro cytotoxicity potency (IC50, nM) of R462-CPT2 and R465 CPT in six CRC cell lines.

| Cell Line | RNA-seq data of LGR4/5/6 expression (in RPKM) |  |  |  | Cytotoxic IC<sub>50</sub> (nM) |  |  |
| --- | --- | --- | --- | --- | --- | --- | --- |
|  | LGR4 | LGR5 | LGR6 | LGR4/5/6 sum | R462-CPT2 | R465-CPT2 | R462-/R465-CPT2 |
| Colo201 | 0.5 | 12.3 | 0.9 | 13.7 | 1.0 | - | - |
| HT29 | 32.9 | 1.8 | 0.3 | 35.0 | 3.3 | 9.1 | 3 |
| Lim1215 | NA | NA | NA | NA | 1.3 | 30.0 | 24 |
| LoVo | 19.8 | 110.9 | 3.1 | 133.8 | 0.9 | 3.1 | 4 |
| LS180 | 17.0 | 28.3 | 1.5 | 46.9 | 0.4 | 9.5 | 26 |
| SW620 | 2.9 | 18.4 | 14.3 | 35.6 | 0.2 | 15.9 | 99 |

Western blot analysis showed that the protein level of LGR4 was largely consistent with LGR4 mRNA level across the six cell lines (Fig. 3a). For LGR5, protein and mRNA levels were also largely consistent except in HT29 cells, with LOVO cells having the highest level of LGR5 protein for its highest level of LGR5 mRNA level (Fig. 3b). R462-CPT2 displayed dose-dependent cytotoxicity with IC50s from 0.16 to 3.3 nM and nearly 100% efficacy except in LS180 cells (Fig. 3c-h). R465-CPT2, on the other hand, displayed IC50s from 3 nM to 30 nM, approximate 3- to 100-fold less potent than R462-CPT2 (Fig. 3c-h) (Table 1).

**Figure 3.**
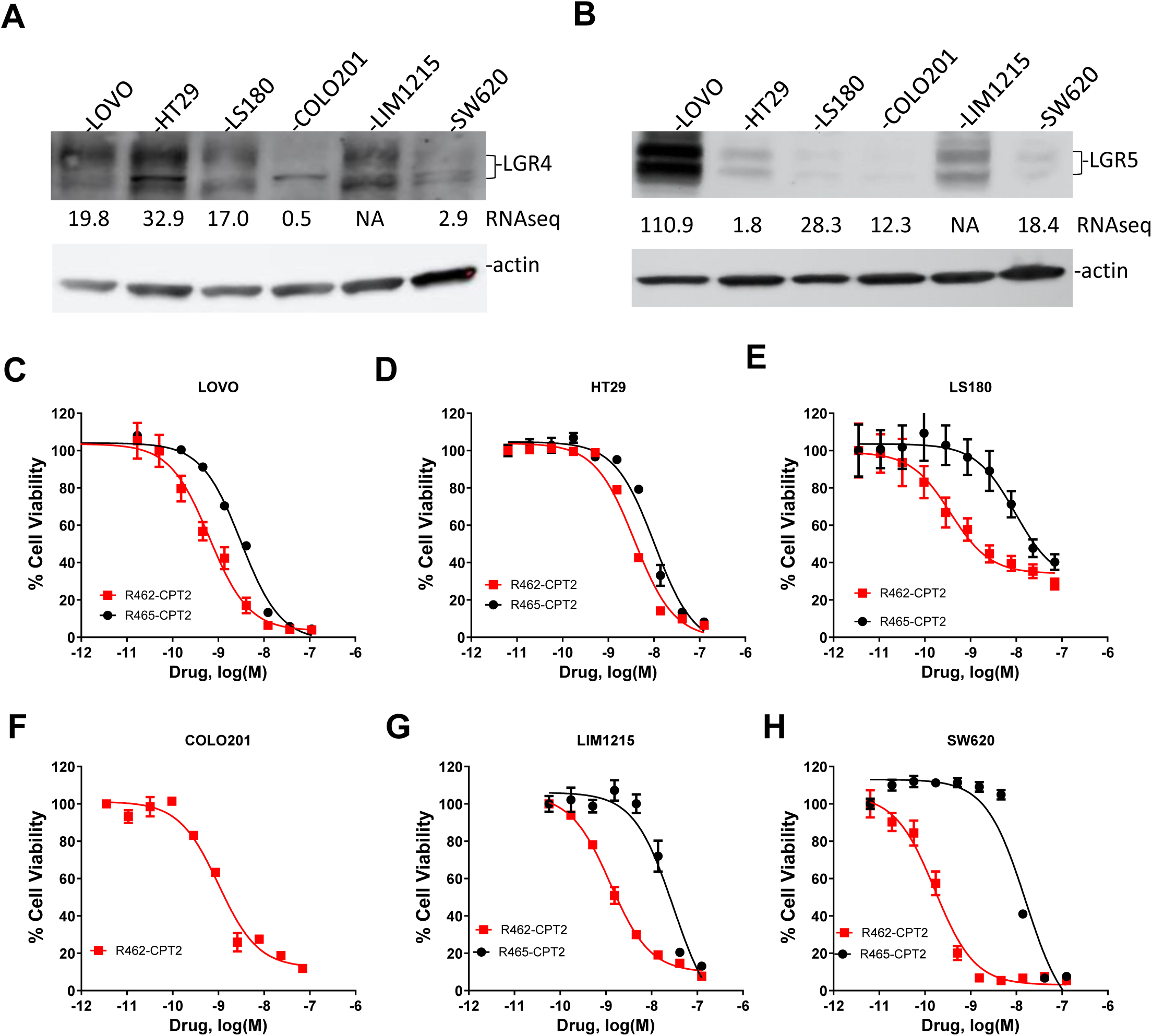
Expression profile of LGR4/5 and In vitro cytotoxicity of R462-CPT2 and R465-CPT2 in six CRC cell lines. **A**, western blot analysis of LGR4 in six CRC cell lines. The numbers underneath each lane denote the values of RNAseq (RPKM) from the CCLE datable. NA, RNAseq data not available in CCLE. **B**, Western blot analysis of LGR5 in the same cell lines. **C-H**, dose-response curves of R462-CPT2 and R465-CPT2 in LOVO cells (**C**), HT29 cells (**D**), LS180 cells (**E**), COLO201 cells (**F**), LIM1215 cells (**G**), and SW620 cells (**H**). The graphs shown here are one representative experiments that were repeated at least once. Error Bars are S.E.M (N= 3).

To understand the mechanism of R465-CPT2-mediated cytotoxicity, we compared R462-CPT2 and R465-CPT2 in HEK293T cells over-expressing LGR4, LGR5, or LGR6 along with vector control cells. R462-CPT2 and R465-CPT2 showed nearly identical potency and efficacy in vector control cells (Supplementary Fig. S2a). In cells overexpressing any of LGR4/5/6, R462-CPT2 was approximately 10-fold more potent than R465-CPT2 (Supplementary Fig. 2b-d).

Together with the binding data of R462-CPT2 and R465-CPT2, these results suggest that conjugation of eight CPT2 molecules introduced the same level of non-specific binding to both R462 and R465, leading the same non-specific potency.

### R462-CPT2 was highly effective in inhibiting tumor growth in vivo

To evaluate anti-tumor activity of R462-CPT2 in vivo, we first determined its stability in normal mice. Previously, we found that R427 conjugated with MMAE had a relatively short half-life (38). R462 and R462-CPT2 were administered at 5 mg/kg and blood were collected at 1, 24, 48, and 72 hours, and peptibody/PDC concentrations in plasma were determined using ligand binding assay on HEK293T cells over-expressing LGR5. Unconjugated and conjugated R462 had relatively short half-life (∼8 hours) (Supplementary Fig. S3), similar to that of R427.

We first compared anti-tumor activity of R462-CPT2 and R465-CPT2 in LoVo cell line xenografts. When tumors reached an average size of ∼150 mm^3^, the animals were randomized into four groups and were given vehicle, R462-CPT2 or R465-CPT2 once every other day. As shown in Figure 4a, R462-CPT2 reduced tumor size by 90% when compared to vehicle (p < 0.05 vs vehicle) while the control PDC R465-PDC2 reduced tumor size by 35% (p > 0.05 vs vehicle). No gross toxicity was observed in any of the treatment groups and no significant changes in body weights were found (Fig. 4b). At termination, all the tumors were excised and weighed. As shown in Fig. 4c, R462-CPT2 and R465-CPT2 treatments led to decreases of 84% and 40%, respectively, in tumor weights when compared to the vehicle group. Importantly, R462-CPT2 showed significantly more tumor weight reduction than R465-CPT2 (p = 0.001, Fig. 4C).

**Figure 4.**
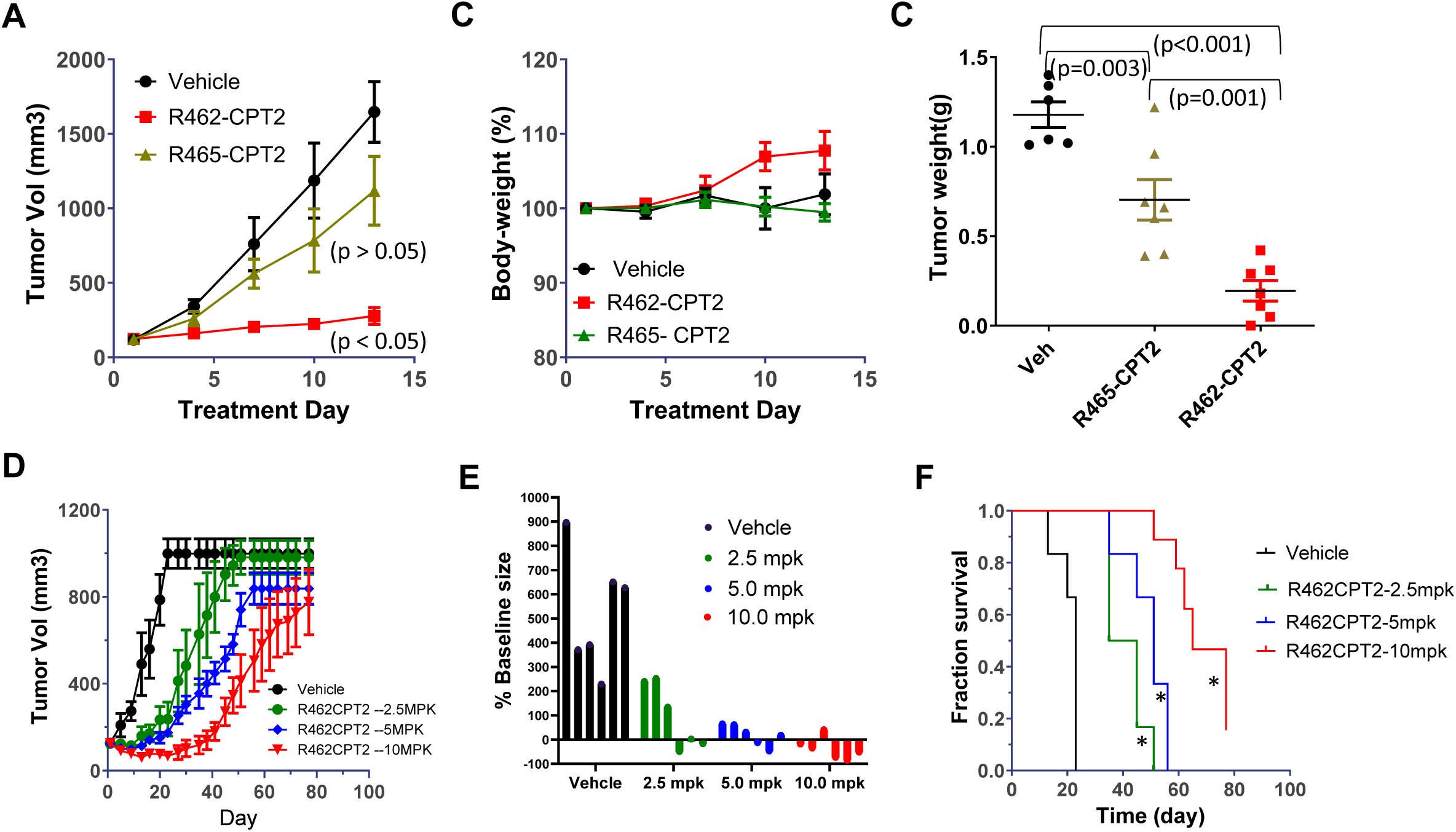
Anti-tumor potency of R462-CPT2 in xenograft model of LOVO cells. **A**, tumor growth curves of LOVO xenografts dosed with vehicle, R462-CPT2, or R465-CPT2 at 5.0 mg/kg, once every other day for a total of eight doses. N = 6-7 per group. Indicated p values are vs vehicle and were calculated by one-way ANOVA followed by Tukey’s multiple comparison test. **B**, Body weight curves of animals in **A**. **C**, tumor weights at termination. Indicated p values were calculated by one-way ANOVA followed by Tukey’s multiple comparison test. **D**, tumor growth curves of LOVO xenografts dosed with R462-CPT2 at 2.5, 5.0, and 10.0 mg/kg, once every other day for a total of eight doses. N = 6 per group. **E**, individual tumor size changes when compared to base line on Day 20. **F**, Kaplan-Meier survival plot of animals doses with R462-CPT2 at three dose levels. *p = 0.001 vs vehicle, Log-rank (Mantel-cox) test.

We then tested anti-tumor potency and efficacy of R462-CPT2 in the LoVo xenograft model using different dose levels. The PDC was given at 2.5, 5.0, and 10.0 mg/kg, once every other day for a total of eight doses when tumors reaches an average size of ∼120 mm^3^. As shown in Figure 4d, R462-CPT2 showed dose-dependent inhibition of tumor growth. At Day 20, five days after the last dose of R462-CPT2 was given, the PDC showed an average inhibition of 83% and 96% in tumor growth at 2.5 mg/kg and 5.0 mg/kg, respectively (Fig. 4e). At 10 mg/kg, five of the six tumors had reduced tumor size with one mouse having no palpable tumor (Fig. 4e). The animals were followed for 11 weeks, and the PDC treatment led to significant increase in overall survival with median survival of 23 days, 40 days (p = 0.001 vs vehicle), 51 days (p = 0.001 vs vehicle), and 65 days (p = 0.001 vs vehicle) for vehicle, 2.5 m/kg, 5.0 mg/kg, and 10.0 mg/kg, respectively, despite that tumors eventually came back in nearly all animals except one in the 10.0 mg/kg group (Fig. 4f). Overall, these results demonstrated that R462-CPT2 had robust and specific anti-tumor effect in a CRC cell line xenograft model.

Next, we evaluated R462-CPT2 in three PDX models (CRC-001 , XSTGI007, and XSTGI010) of CRC (45). The three models express LGR4 and LGR5 at moderate levels with all models being MSS. First, we compared R462-CPT2 and the control PDC R465-CPT2 in the model XSTGI010, and found that R462-CPT2 inhibited tumor growth by 82% whereas the control PDC R465-CPT2 only inhibited tumor growth by 43%, which was significantly different (p < 0.0001, one-way ANOVA followed by Tukey multiple comparison test), suggesting that the anti-tumor activity of R462-CPT2 was largely specific. In the models CRC-001 and XSTGI007, R462-CPT2 was able to significantly inhibit tumor growth by 80% and 83% (p < 0.0001 vs vehicle, T test, for both models), respectively (Fig. 5c-f). In all studies, no gross toxicity was observed with no significant body weight change observed in any of the groups. Overall, these results indicated that R462-CPT2 was able to inhibit tumor growth by approximately 80% following dosing at 5 mg/kg, every other day, and this activity was largely mediated by targeting LGR4/5/6 specifically.

**Figure 5.**
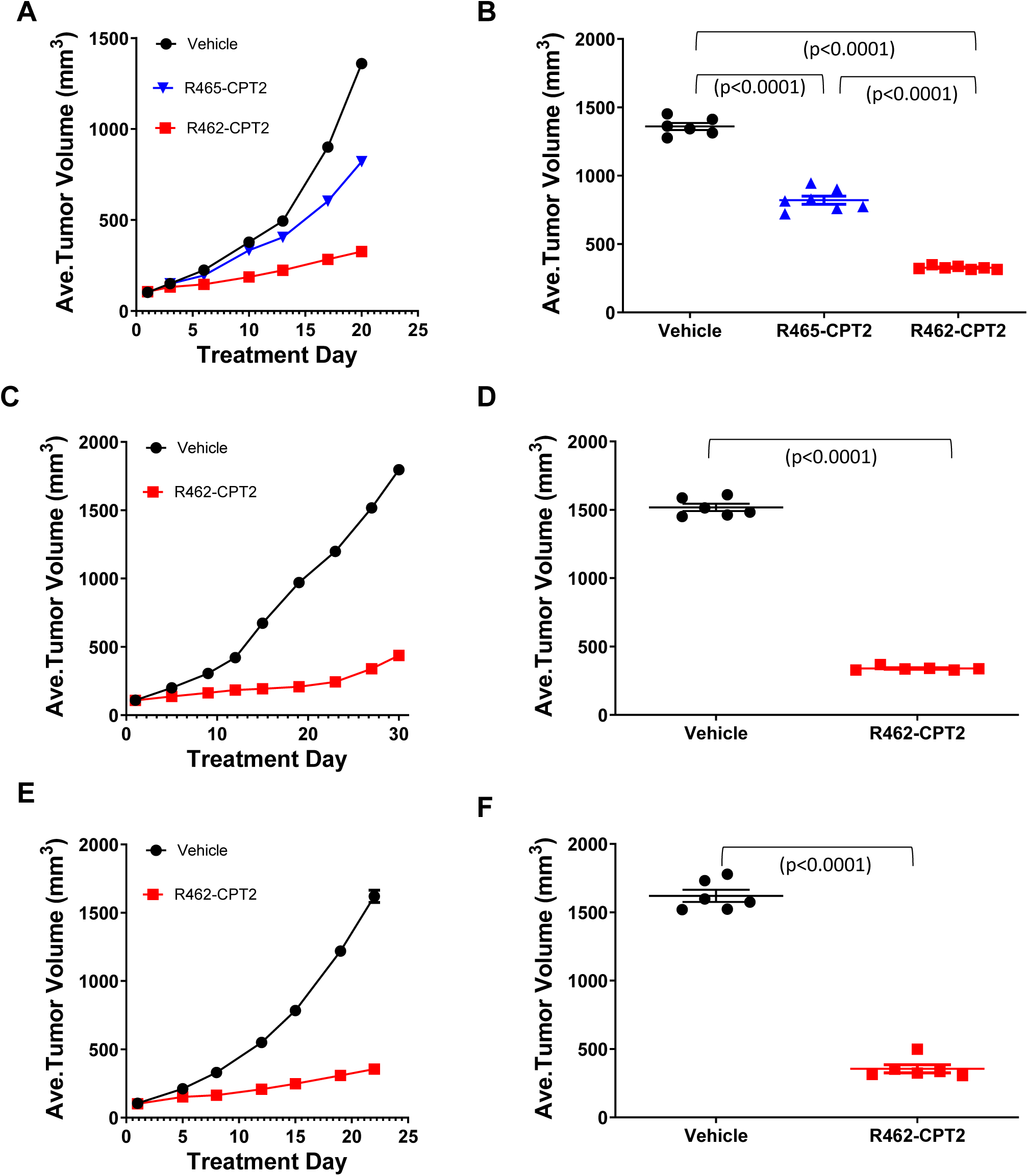
Anti-tumor effect of R462-CPT2 in three PDX models of CRC. **A**, tumor growth curves of PDX model XSTGI010 dosed with R462-CPT2, R465-CPT2 at 5.0 mg/kg, or vehicle alone. **B**, Individual tumor size on Day 20 of A. **C**, tumor growth curves of PDX model CRC-001 dose with R462-CPT2 at 5.0 mg/kg or vehicle alone. **D**, individual tumor size of animals in C on day 30. **E**, tumor growth curves of PDX model XSTGI007 dose with R462-CPT2 at 5.0 mg/kg or vehicle alone. **F**, individual tumor size of animals in **E** on day 22.

### R462-CPT2 had no major adverse effect at therapeutically effective dose levels

Given the robust anti-tumor efficacy of R462-CPT2 in cell line xenograft and PDX models, we carried out a comprehensive toxicity study in normal mice. The PDC was given at 5.0, 10, and 15 mg/kg, along with vehicle control group to C57Bl/6 mice (3M/3F per group), every other day for a total of three doses by IP injection. The animals were monitored for general well-being and body weight. At termination on Day 8, animals were weighted and blood was collected for complete blood count and serum chemistry analyses. Animals were necropsied with weighing of 7 organs and collection of 25 tissues for histopathology analysis. Tissues were processed routinely for histology and examined by a board-certified veterinary pathologist.

In a toxicity test in normal mice, no major adverse effect in any of the treatment groups was found in body weights, serum chemistry, and blood count, except the decrease in red blood cells which were still in the normal range (Fig. 6). In the hematopoietic system, the medium and high dose groups (10 and 15 mg/kg) showed mild reduced cellularity in the thymus and spleen, and increased spleen weight was observed in the 15 mg/kg group. Hematopoietic toxicity is a common side effect CPT-based ADC therapy (41). Interestingly, in the paper by Lyski et al (2), a CPT2-based ADC with the same PEG8-VKG-linker was well tolerated at up to 60 mg/kg in normal rats. Since ADC-based toxicity is mostly due to non-specific release of payloads (50), these results suggest that CPT2-based ADCs/PDCs may be tolerated at high dose levels. In addition, the finding that no adverse effect was observed on the gastrointestinal system even at the highest dose level strongly suggest that simultaneous targeting of LGR4/5/6 by RSPO4 furin domain-based PDC is a viable approach despite the expression of LGR5 in adult stem cells of the intestine. The lack of toxicity in the GI system is most likely due to the relative low expression of LGR4/5/6 in normal stem cells when compared to the cancer cells.

**Figure 6.**
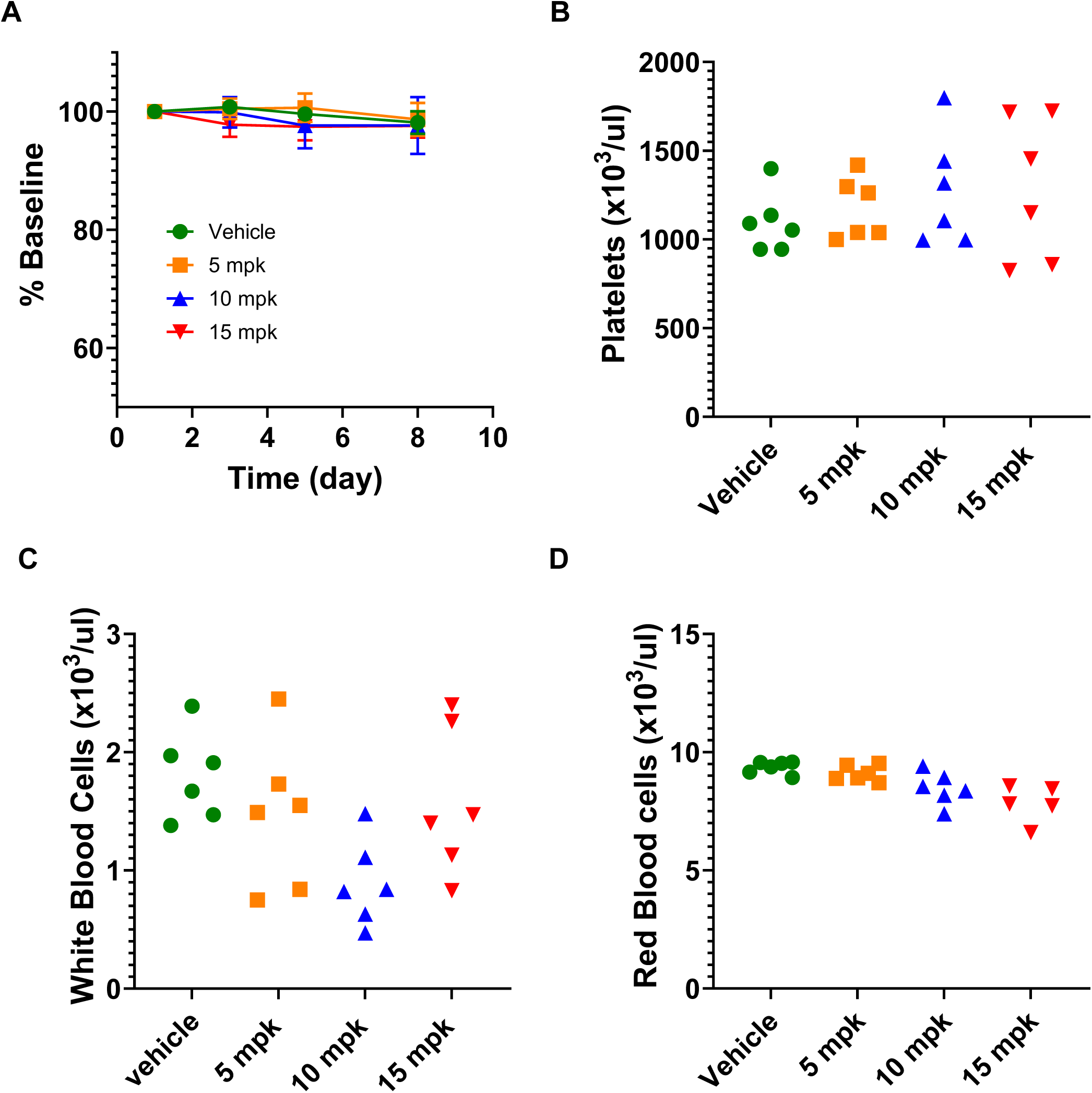
Mouse toxicity study results. **A**, body weight normalized to baseline of each animal. **B**, Platelet counts. **C**, white blood cell count. **D**. Red blood cell count.

## Discussion

LGR4/5/6 have been shown to be expressed at moderate to high levels in CRCs by multiple independent studies and anti-LGR5 ADCs demonstrated potential as a potential treatment of CRC (27,51). As not all CRC cells express LGR5 even in tumors with high overall of LGR5 expression and LGR5-positive and LGR5-negative cell interconvert (52), we set out to develop a strategy that utilizes a mutant ligand of LGR4/5/6 that retains high affinity receptor binding without potentiating Wnt/β-catenin signaling (38). Here we generated a new peptibody (R462) of mutant RSPO4 furin domain and conjugated with an analog of camptothecin called CPT2 that is more potent and soluble than the payload used in the FDA-approved ADCs (41).

Camptothecin-derived payloads are now the leading class for ADC development due to its success in solid tumors, and irinotecan, a camptothecin derivative, is an effective chemotherapy against CRC. R462-CPT2 showed potent cytotoxic activity in vitro in multiple CRC cell lines with LGR4/5/6 expression at various level and strong anti-tumor effect in vivo in cell line xenograft and PDX tumors.

By building upon previous work with R427, we engineered R462 with Q295 and Q297 residues in the Fc domain to enable conjugation of up to eight CPT2 molecules per peptibody, for enhanced conjugation capacity. The fusion of RSPO4 furin domain to IgG1-Fc not only extended its half-life in vivo but also increased its binding affinity toward LGR4 due to the dimeric nature of the Fc domain. We also created a control peptibody R465 that is identical to R462 except that it is unable to bind to LGR4/5/6 due to two amino acid changes in the Fu2 domain of RSPO4 furin domain. R462 showed highly potent and specific binding to LGR4 and LGR5 with no binding to LGR4/5-negative cells, indicating that the peptibody is highly specific. As expected, R465 showed no binding to LGR4/5-postive or -negative cells. However, conjugation of eight molecules of CPT2 to R465 using click chemistry led to a low affinity, non-specific binding to cells with or without over-expressing LGR4/5, indicating that conjugation of 8 molecules of CPT2 led to non-specific binding, even with a PEG8 linker. Previously, we had 4 molecules of the payload mono-methyl auristatin (MMAE) conjugated to a similar RSPO4 peptide without observing non-specific binding. Since MMAE is more soluble and only 4 molecules were conjugated per peptibody, the low affinity, non-specific binding of R465-CPT2 is most likely due to the hydrophobic nature of CPT analogs (40). The non-specific activity of the PDC could be also due to aggregation of CPT2 because of the close proximity of the two Gln residues used for conjugation. Future work will explore the use of Gln residues that are located far apart on the Fc domain.

In the LoVo cell line xenograft model, R462-CPT2 demonstrated significantly more efficacious anti-tumor effect than R465-CPT2 (Fig. 4A), indicating that R462-CPT2 was able to deliver the payload more effectively to the cancer cells. The level of anti-tumor activity of R465-CPT2 could be due to non-specific release of free payload in circulation or in tissues, which is known to contribute to anti-tumor efficacy of ADCs (53,54). Furthermore, R462-CPT2 showed dose-dependent anti-tumor effect in the LoVo xenograft model with 4/6 tumors in regression and 1/6 in complete remission (Fig. 4D-E). However, tumors eventually grew back after treatment stopped, albeit at a slow rate (Fig. 4D), suggesting that continuous treatment was necessary to suppress tumor growth. In the CRC PDX models, R462-CPT2 only showed ∼80% reduction in tumor growth at 5 mg/kg. High dosage or combination therapy may be required to cause tumor regression or complete remission. Future studies will also evaluate the combination of R462-CPT2 with standard chemotherapy of colon cancer such as FoxFox (folinic acid, fluorouracil and oxaliplatin)(2).

Notably, the anti-tumor activity of R462-CPT2 was achieved with minimal toxicity, even at the highest dose tested (15 mg/kg). Interestingly, despite the expression of LGR5 in adult intestinal stem cells, no adverse effects were observed in the gastrointestinal system. This finding suggests that the relatively low expression levels of LGR4/5/6 in normal stem cells compared to cancer cells may provide a therapeutic window for targeting these receptors with PDCs. However, further studies are needed to fully understand the potential for on-target but off-tumor toxicities.

In conclusion, we generated a new peptibody based on a mutant RSPO4 furin domain that showed high affinity, specific binding to LGR4/5/6, and its drug conjugate displayed potent cytotoxic activity in CRC cell lines expressing any of LGR4/5/6 in vitro, and robust anti-tumor efficacy in vivo without major adverse effect. Future studies will focus on optimizing the in vivo stability of the peptibody and identify strategies that increase the solubility of the PDC.

## Acknowledgements

The authors would like to thank Mr. Jack Adams of UTHealth-Houston for drawing the structure of PEG2-bis-PE3-azido-DBCO-PEG8-VKG-CPT2. This work was supported in part by funding from the Cancer Prevention and Research Institute of Texas (CPRIT) RP220169 (to Q.J.L. and K.S.C.), RP210119 (to Q.J.L.), National Cancer Institute (NCI) R01CA226894 and R21 CA282378 (to K.S.C), and the Janice David Gordon for Bowel Cancer Research Endowment (to Q.J.L.).

