## Supplementary Figures for "Anti-tumor activity of camptothecin analog conjugate of a RSPO4-based peptibody targeting LGR4/5/6 in preclinical models of colorectal cancer"

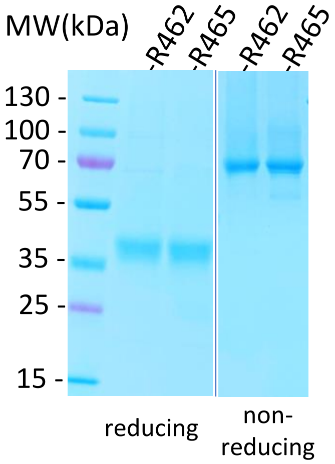


Supplementary Figure S1. Coomassie blue staining image of purified R462 and R465 under reducing (left side) and non-reduction condition.

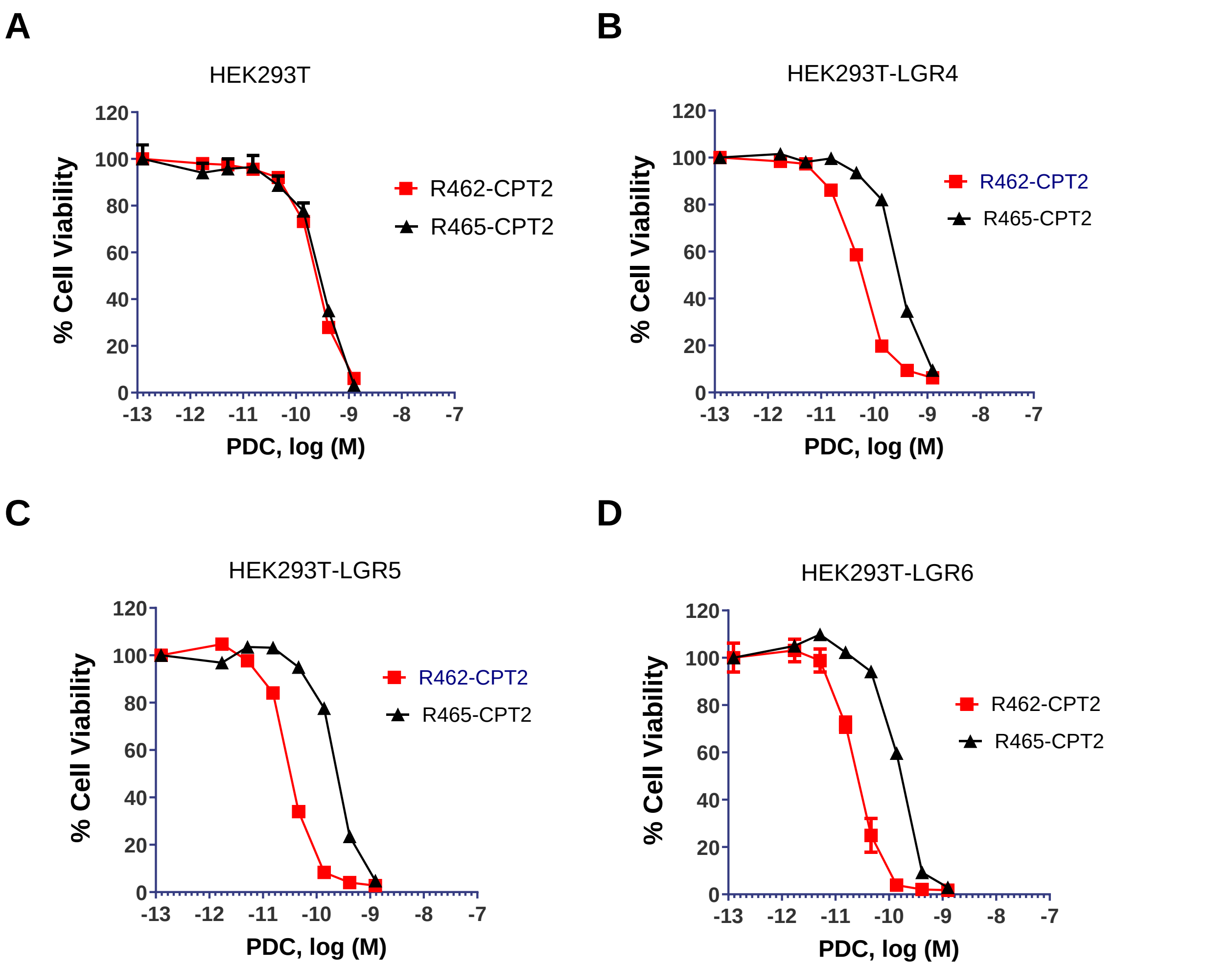


Supplementary Figure S2. In vitro cytotoxicity of R462-CPT2 and R465-CPT2 in HEK293T cells and HEK293T cells expressing LGR4, LGR5, or LGR6. **A**, HEK293T parental cells. **B**, HEK293T-LGR4 cells. **C**, HEK293T-LGR5 cells. **D**, HEK293T-LGR6 cells. Graphs are one representative of experiments that were repeated at least once. Error bars are S.E.M (N= 3).

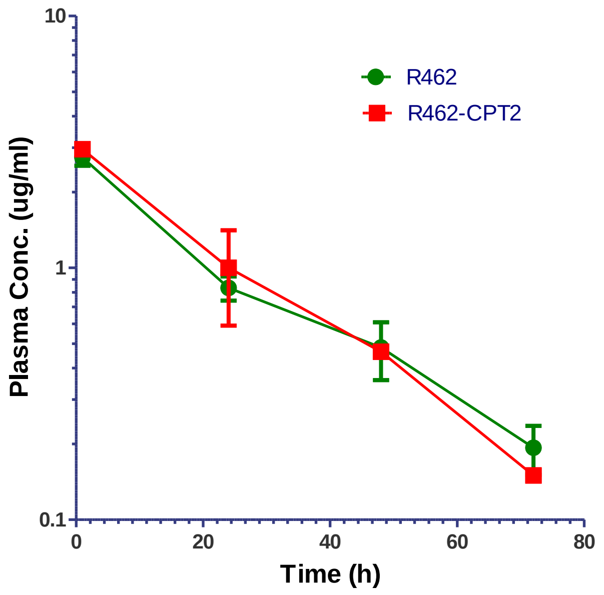


Supplementary Figure S3. Pharmacokinetics of R462 and R462-CPT in mice. R462 and R462-CPT2 were injected into mice at 5.0 mg/kg by intraperitoneal administration, and blood was collected at the indicated time points. Peptibody/PDC concentration were determined by receptor binding assays using HEK293T-cells stably expressing LGR5.
